# Functional transcriptomics in diverse intestinal epithelial cell types reveals robust gut microbial sensitivity of microRNAs in intestinal stem cells

**DOI:** 10.1101/087882

**Authors:** Bailey C. E. Peck, Amanda T. Mah, Wendy A. Pitman, Shengli Ding, P. Kay Lund, Praveen Sethupathy

## Abstract

Gut microbiota play an important role in regulating the development of the host immune system, metabolic rate, and at times, disease pathogenesis. The factors and mechanisms that mediate communication between microbiota and the intestinal epithelium are poorly understood. We provide novel evidence that microbiota may control intestinal epithelial stem cell (IESC) proliferation in part through microRNAs (miRNAs). We demonstrate that miRNA profiles differ dramatically across functionally distinct cell types of the mouse jejunal intestinal epithelium and that miRNAs respond to microbiota in a highly cell-type specific manner. Importantly, we also show that miRNAs in IESCs are more prominently regulated by microbiota compared to miRNAs in any other intestinal epithelial cell (IEC) subtype. We identify miR-375 as one miRNA that is significantly suppressed by the presence of microbiota in IESCs. Using a novel method to knockdown gene and miRNA expression *ex vivo* enteroids, we demonstrate that we can knockdown gene expression in Lgr5^+^ IESCs. Furthermore, when we knockdown miR-375 in IESCs, we observe significantly increased proliferative capacity. Understanding the mechanisms by which microbiota regulate miRNA expression in IESCs and other IEC subtypes will elucidate a critical molecular network that controls intestinal homeostasis and, given the heightened interest in miRNA-based therapies, may offer novel therapeutic strategies in the treatment of gastrointestinal diseases associated with altered IESC function.

---

The intestinal epithelium is a single layer of cells exposed to the intestinal lumen, and is composed of multiple cell types including the proliferative IESCs and progenitor cells (also known as transit amplifying cells), as well as differentiated absorptive enterocytes and secretory goblet, Paneth, and enteroendocrine cells (EECs). IESCs divide to yield more rapidly proliferating progenitors that give rise to all of the other IEC types and drive continuous renewal of the intestinal epithelium every ~3-5 days(1). Proper renewal facilitates important intestinal epithelial functions including barrier integrity to protect against invasion of harmful toxins present in the intestinal lumen, nutrient digestion and absorption, and production of hormones that regulate systemic energy homeostasis. These physiological processes are mediated in part by interactions with resident microbiota(2). Studies using germ-free animals have demonstrated that gut microbiota influence intestinal barrier function, nutrient absorption, proliferation, differentiation, cellular signaling, and migration(3, 4). However, the molecular factors and mechanisms underlying microbiota-mediated control of IEC functions, particularly IESC proliferation, are unknown.

miRNAs have emerged as critical regulatory factors of many biological processes in numerous tissues and are known to confer phenotypic robustness in response to environmental stimuli(5). However, less is known about miRNA expression and function in the intestinal epithelium compared to most other tissues. Recently, miRNAs were implicated in the regulation of IEC physiology(6, 7). McKenna et al. (2010) demonstrated in mice that the IEC-specific knockout of *Dicer1*, an essential enzyme for canonical miRNA biogenesis, results in altered IEC proliferation, differentiation, nutrient absorption, and impaired barrier function, indicating that miRNAs are likely important modulators of intestinal homeostasis(6). Furthermore, the presence of microbiota in the gut has been shown to alter miRNA expression profiles in intestinal macrophages(8), as well as in whole intestine (9, 10). Understanding the mechanisms by which microbiota regulate miRNA and gene expression in IESCs and other IEC subtypes will elucidate a critical molecular network that controls intestinal homeostasis and, given the heightened interest in miRNA-based therapies, may offer novel therapeutic strategies in the treatment of gastrointestinal diseases associated with altered IESC function. However, no study to date has investigated miRNA expression and activity across the functionally distinct IEC subtypes, and cell-type specific effects of microbiota on miRNAs is completely unknown. We hypothesized that each IEC population has a distinct miRNA profile, and that miRNAs respond to gut microbiota in a cell-type specific manner in order to control function and overall homeostasis of the intestinal epithelium.

## RESULTS

*Germ-free mice have an altered jejunal IEC composition compared to their conventionalized and conventionally-raised counterparts*—We selected the well-characterized Sox9-EGFP transgenic mouse model to evaluate miRNA expression and response to microbiota in functionally distinct IECs. This model was originally created by GENSAT(11), who developed the model by randomly inserting into the mouse genome a BAC containing the EGFP gene driven by the cloned genomic regions upstream and downstream of *Sox9*(12). A beneficial feature of this model is that EGFP expression is fully penetrant within the mouse intestine, which permits the isolation and analysis of four distinct IEC populations(12). Applying the same fluorescence-activated cell sorting (FACS)-based approach used to isolate IESCs from the commonly used Lgr5-EGFP model, which demonstrates mosaic expression among crypts in the intestine, both actively cycling IESCs (Sox9^^Low^^) and transit-amplifying progenitor cells (Sox9^Sublow^) can be isolated from the Sox9-EGFP mouse intestine(12, 13). Moreover, two additional differentiated cell populations can also be isolated on the basis of variable EGFP intensity, including Sox9^Neg^ (mostly differentiated enterocytes as well as goblet cells and Paneth cells), and Sox9^High^ (primarily EECs as well as reserve/quiescent +4 stem cells) (12-17).

To evaluate the effect of microbiota on miRNA expression in IECs, we first took a conventionalization approach (Figure 1a). A two-week conventionalization was selected because previous studies show this to be a time point at which the gene expression profile begins stabilizing in the small intestine of young mice following conventionalization(18-20). After generating germ-free (GF) Sox9-EGFP animals at the University of North Carolina at Chapel Hill (UNC) Gnotobiotic core facility, we selected four pairs of female GF Sox9-EGFP littermates from 4 different litters born between February and July 2015. One littermate from each pair was randomly selected at 8-10 weeks of age for conventionalization. The 2-week conventionalization resulted in slightly decreased body weight relative to the remaining germ-free sibling, along with a commensurate increase in liver weight (Supplemental Figure 1). However, no differences were observed in length of the small intestine or colon between GF and conventionalized (CV) animals (Supplemental Figure 1). IECs were collected from the mid-region of the small intestine (Methods), hereafter referred to as jejunum, of the GF and CV animals and FACS was performed based on Sox9-EGFP intensity (Figure 1a). Special care was taken to gate out cellular debris, dead and dying cells, immune cells, and multiplets during FACS (See Methods, Supplemental Figure 2). Additionally, a strict gating scheme was used to avoid contamination between cell populations.

**FIGURE 1.**
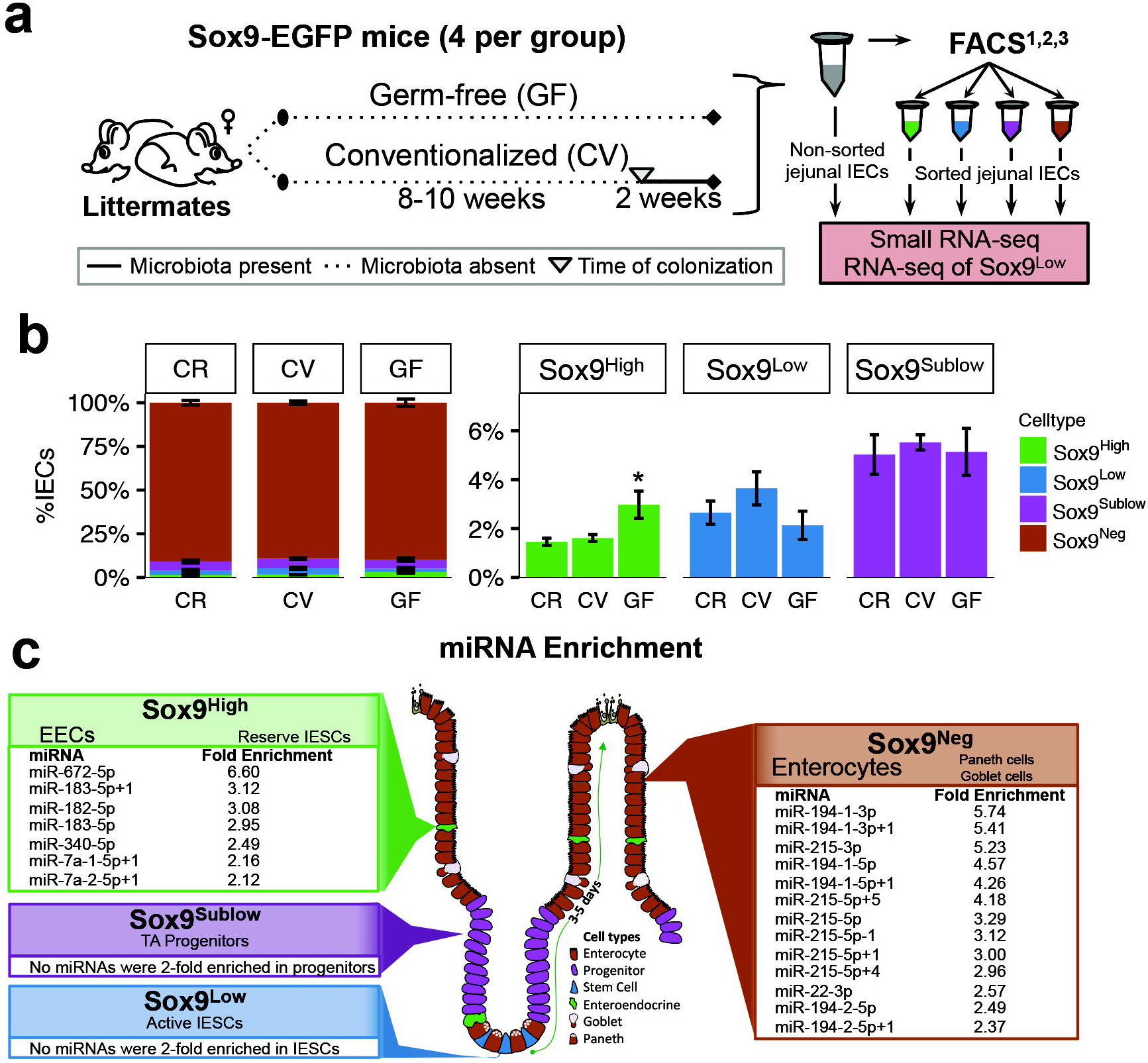
The Sox9-EGFP mouse model for characterizing subpopulations of the mouse intestinal epithelium. **a**) Diagram of our experimental design to profile the transcriptional landscape of distinct intestinal epithelial cell (IEC) populations and their response to microbial conventionalization. **b**) Mean percentage of each IEC subtype sorted from jejunum of conventionally-raised (CR), germ-free (GF) and conventionalized (CV) mice (n=4 each). Error bars depict standard error of the mean. Significance determined using two-tailed unpaired t-Test relative to CR, and is denoted as follows: * p < 0.05. **c**) Cartoon showing location and types of IECs in the Sox9-EGFP animal. Listed are miRNAs with mean expression in reads per million mapped to miRNAs (RPMMM) across all CR, CV, and GF animals that were found to be at least 2-fold enriched in each respective population. Fold enrichment indicates enrichment of the miRNA to the IEC subpopulation with the next highest mean expression. Only miRNAs with RPMMM > 400 in at least one sample are included in the analysis.

**FIGURE 2.**
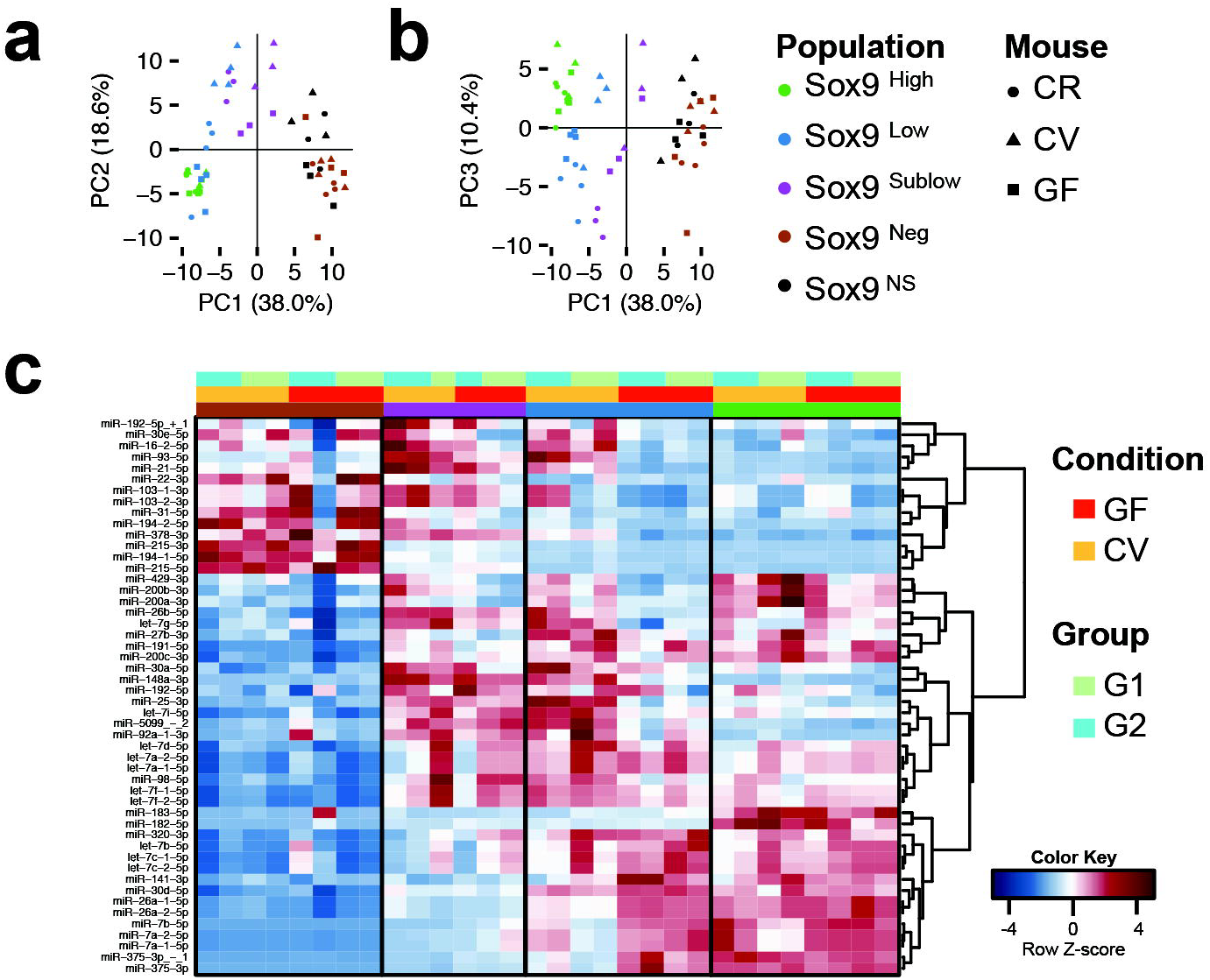
miRNAs in the intestinal epithelium show cell type specific expression and responses to microbiota. **a & b**) Principal components analysis (PCA) applied to miRNA expression in reads per million mapped to miRNAs (RPMMM) for all miRNAs that reached an expression threshold of at least 400 in 1+ samples (n=149). Biplots of variance projections of PC1 on PC2 (**a**) and PC1 on PC3 (**b**) are shown. Percent variance explained by each principal component is shown in parentheses along each axis. Samples are colored based on the cell population, and the shape of the point indicates whether the sample came from a conventionally-raised (CR), germ-free (GF) or conventionalized (CV) mouse. **c**) Hierarchical clustering of the top 50 most highly expressed miRNAs across sorted intestinal epithelial cell populations across all categories of mice (GF, CV, and CR). Color bars denote cell population, condition, and sequencing group (G1 or G2).

We began by comparing the abundances of each major IEC subpopulation in GF and CV animals to conventionally-raised (CR) chow-fed animals. We found very similar abundances of Sox9^Sublow^ cells (transit amplifying) and Sox9^Neg^ cells (enterocytes, Paneth, and goblet) in CV and GF populations relative to CR mice (Figure 1b). Notably, GF mice had significantly more Sox9^High^ cells (EECs) than CR mice (fold change=2.04, p=0.04, Figure 1b), which is consistent with previous studies comparing EECs in the jejunum of GF and CR rodents(21, 22). Also, there were on average fewer Sox9^Low^ cells (actively cycling IESCs) in GF mice relative to CR and CV mice, which has not been shown before, but could help explain previous reports suggesting reduced proliferation in the small intestine of GF animals(23-25).

*IESCs demonstrate robust transcriptional changes in response to gut microbiota*—To evaluate the transcriptional changes that occur in response to conventionalization in the IESCs of GF and CV animals, we performed RNA-sequencing analysis on the Sox9^Low^ population (which we will refer to as IESCs for simplicity). We identified 823 genes and long, non-coding RNAs (lncRNAs) significantly elevated in GF IESCs and 334 genes and lncRNAs significantly elevated in CV IESCs (Supplemental Figure 3a). Gene Ontology Biological Process(26, 27) enrichment analysis using Enrichr(28) revealed that genes elevated in CV IESCs are most significantly over-represented in pathways related to proliferation such as ‘mitotic cell cycle’ and ‘nuclear division’ (Supplemental Figure 3b). The genes elevated in GF IESCs genes were associated with processes related to hormone secretion and transport (Supplemental Figure 3b). Consistent with these findings, we observed that established markers of proliferation (*Ccnb1*, *Cdk1*, and *Mki67*) are significantly up-regulated, positive transcriptional regulators of IESC proliferation and self-renewal (*Gata4* and *Gata6*) are up-regulated, and negative regulators of IESC proliferation and self-renewal (*Bmp4*) are down-regulated in CV IESCs (Supplemental Figure 3c). Also, some, but not all, classic markers of enteroendocrine cells are upregulated in GF IESCs (Supplemental Figure 3c), which could indicate some priming for cells to enter the EEC lineage, consistent with our observation that GF mice have more Sox9^High^ cells. Known markers of reserve (quiescent) stem cells were not significantly different between CV and GF IESCs (Supplemental Figure 3c), nor were markers for Paneth cells (*Lyz*), goblet cells (*Muc2*), and enterocytes (*Elf3*). These data confirm that the Sox9^Low^ cells are indeed enriched for IESCs and that CV IESCs harbor a gene signature consistent with increased proliferative capacity. As miRNAs are known regulators of proliferation and differentiation, we performed small RNA-sequencing of each of the functionally distinct IEC subpopulations from four CR, GF, and CV animals.

**FIGURE 3.**
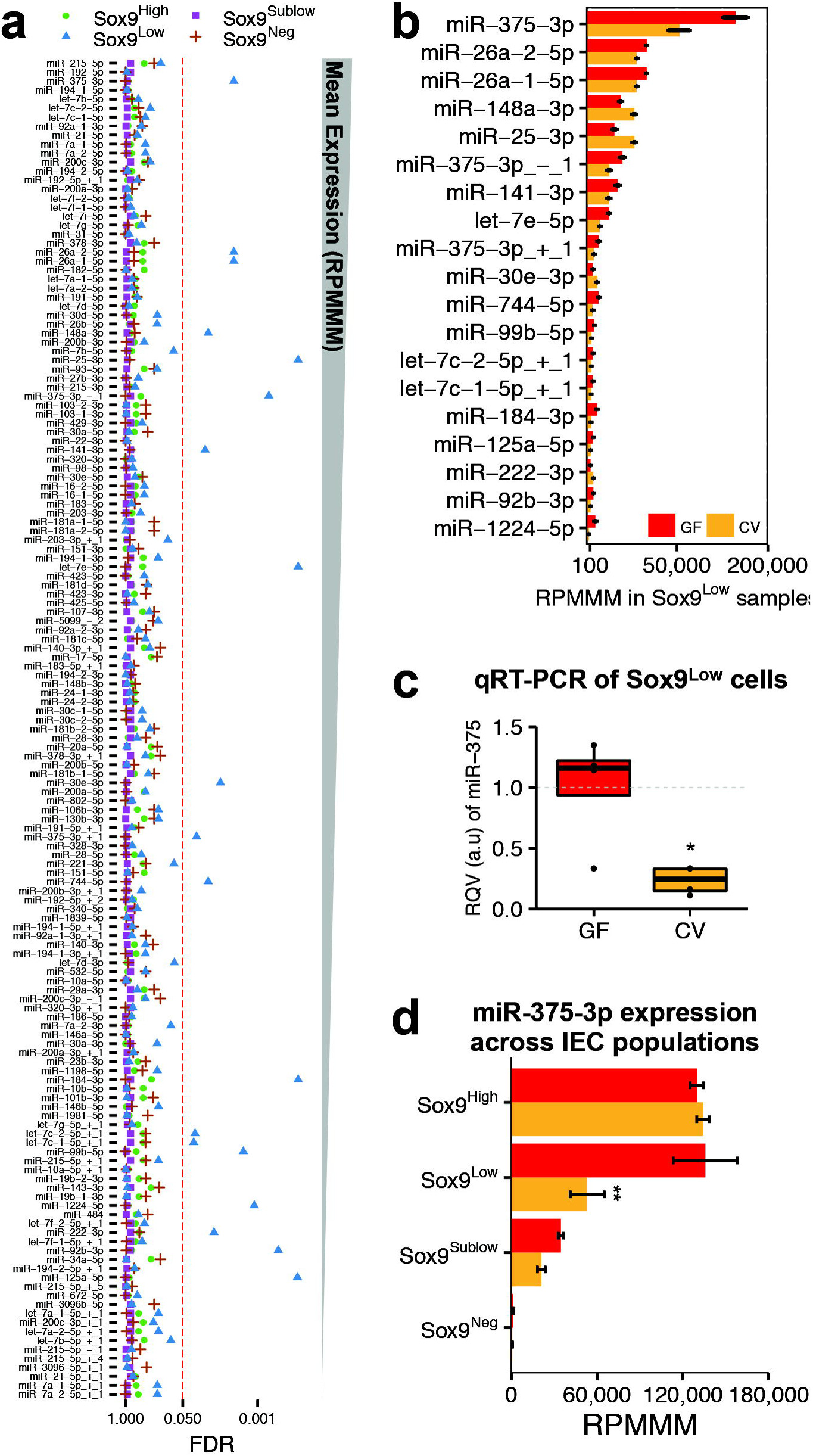
Cell population specific response of miRNAs to conventionalization revealed through linear modeling analysis. **a**) A linear regression model was used to account for the following covariates: cell population, condition, sequencing batch, and littermate pair. For each miRNA that met an expression threshold of reads per million mapped (RPMMM) > 400 in 1+ samples, the False Discovery Rate (FDR) multiple testing correction of the cell population*condition covariate interaction p-value is plotted. miRNAs are ordered by average expression across all intestinal epithelial cell (IEC) subtypes across all categories (GF, CV, and CR), and vertical red dashed line indicates FDR = 0.05. Cell population is signified by color and shape. **b**) The mean expression in RPMMM of the 19 miRNAs identified as having a significant response (FDR < 0.05) to microbiota the Sox9^Low^ intestinal epithelial stem cell population (in panel **a**) are plotted for both germ-free (GF) and conventionalized (CV) mice. X-axis is shown on a square root scale. Error bars depict standard error of the mean. **c**) qRT-PCR confirming miR-375 is reduced upon conventionalization (n=4 for each condition). Data are shown in a standard box-and-whisker plot with median displayed as thick horizontal line, shaded region depicting the inner quartile range (IQR), and whiskers extending to the maximum and minimum data points that fall within 1.5*IQR. * p < 0.05, two-tailed Student’s t-Test. **d**) Mean expression of miR-375 in each cell population are shown in RPMMM for both GF and CV mice. Error bars depict standard error of the mean. Significance determined as indicated in panel **a**.

*miRNAs show cell-type specific expression across functionally distinct populations of IECs*—Total RNA was isolated from the four sorted populations from each animal, as well as from non-sorted IECs (NS IECs; NS IECs were purified by FACS, but not sorted based on Sox9-EGFP intensity). Small RNA-sequencing was performed in three batches, two of which contained small RNA libraries from sorted and unsorted IECs from two GF animals and two CV animals. The third batch contained libraries of the four CR animals. miRNAs and their isomiRs were aligned and quantified using miRquant, our previously described method (see Methods for details)(29). To test our hypothesis that miRNAs are differentially expressed among functionally distinct IEC subtypes, we evaluated miRNAs with an expression level of at least 400 reads per million mapped to miRNAs (RPMMM) in one or more samples, identifying 149 robustly expressed miRNAs across all IEC populations.

Many miRNAs were uniquely enriched in one IEC subtype relative to all others (>2-fold more highly expressed than any other cell type across all samples; Figure 1c). For example, we found that miR-215 and miR-194 are enriched in Sox9^Neg^ cells, which consist primarily of enterocytes. Both of these miRNAs are processed from a single primary miRNA transcript on Chr1 and were previously shown to be induced by HNF4α during differentiation of Caco-2 colon carcinoma cells(30). Five miRNAs are enriched in Sox9^High^ cells (EECs and reserve stem cells), including miR-182-5p and miR-183-5p (Figure 1c), which are also generated from a single primary miRNA transcript. Consistent with enrichment in a subpopulation of cells composed largely of EECs, miR-182 has been shown to have important functions in other endocrine cells, specifically, pancreatic beta cells(31). Unexpectedly, we did not find any miRNAs enriched in the Sox9^Low^ IESCs or Sox9^Sublow^ progenitors, which are the only actively proliferating cell populations (Figure 1c).

To further assess variability within and across samples, we performed principal component analysis (PCA) to reduce dimensionality and assess the effect of microbial presence on each sample. The first three principal components (PC1-PC3) captured 67% of the variability across samples. Using biplots we evaluated segregation of samples by cell type and mouse condition (GF, CV, or CR). We showed that miRNA expression profiles are sufficient to cluster most samples by their respective cell types regardless of microbial status (Figure 2a & 2b). For example, Sox9^Neg^ cells and NS IECs are tightly clustered, which is expected given that NS IECs are composed of 85-90% Sox9^Neg^ cells. In the first biplot, GF Sox9^Low^ IESCs do not cluster together with CV and CR Sox9^Low^ IESCs (Figure 2a), which indicates that IESCs are particularly sensitive to the presence or absence of microbiota. However, when PC1 and PC3 are projected, clear cell type specific clustering is observed regardless of mouse condition (Figure 2b), suggesting that the subset of miRNAs loaded into PC2, some 40 miRNAs, contribute to the grouping observed in Figure 2a. Taken together, these data indicate strong cell type specific expression of miRNAs across IEC populations and a robust effect of microbial presence on the IESC population specifically. Both observations are supported by visualization of the most highly expressed miRNAs across GF and CV samples (Figure 2c). *miRNAs of IESCs are the more responsive to microbial presence than other IEC types*—To evaluate the cell type specific responses to microbiota and account for batch and littermate effects, we used a linear modeling approach (See Methods). We found the expression levels of 11 miRNAs (miR-34a-5p, miR-200c-3p, miR-200c-3p-1, miR-143-3p, miR-130b-3p, miR-140-3p+1, miR-378-3p+1, miR-20a-5p, miR-17-5p, miR-93-5p, and miR-29a-3p) to be significantly influenced by microbial status across all cell populations. These miRNAs in general are elevated in CV mice across all cell populations, though the magnitude of effect is always most pronounced in the Sox9^Low^ (IESC) population. When we considered cell type specific responses to microbiota in the functionally distinct subpopulations, we were surprised to find miRNAs are only significantly changing in the IESC population in response to microbiota, whereas no miRNAs were identified as significant changed in the Sox9^High^, Sox9^Sublow^, or Sox9^Neg^ populations (Figure 3a, Supplementary File 1). A total of 19 miRNAs were significantly altered by microbiota in IESCs (Figure 3b), which underscores the highly cell-type specific miRNA response to microbiota.

Of these 19 microbiota-sensitive miRNAs in IESCs, miR-375-3p is ~2.5-fold (FDR=0.003) reduced in CV IESCs compared to GF IESCs and is the most highly expressed (Figure 3a & 3b). Notably, miR-375-3p is 2.4- and 7.2-fold more highly expressed than the next-most significant microbiota-sensitive miRNA in the CV and GF IESC populations, respectively (Figure 3b). We also found that its isomiRs, miR-375-3p-1 and miR-375-3p+1, are also both significantly downregulated in IESCs upon conventionalization (FC=-2.45, FDR=0.0006; and FC=-2.47, FDR=0.02, respectively; Figure 3b). qRT-PCR in Sox9^Low^ cells confirmed that the miR-375-3p family is significantly downregulated by conventionalization (FC= −3.85, p=0.03; Figure 3c). Of note, miR-375-3p exhibits rather low expression in Sox9^Sublow^ progenitors and Sox9^Neg^ cells (Figure 3d). Although miR-375-3p is highly expressed in Sox9^High^ EECs, it is not altered in EECs by conventionalization, and is only significantly downregulated by microbiota in the IESCs.

*Knockdown of gene expression in IESCs of ex vivo enteroids using gymnosis*—To functionally evaluate the effect of the observed miRNA and gene expression changes, we sought out methods to downregulate gene expression in IESCs of *ex vivo* enteroid culture systems, which have been shown to maintain *in vivo* cellular composition and molecular gene expression profiles over time(32). We evaluated the use of gymnosis, a term coined by the Troels Koch laboratory in 2009(33), to describe a process of introducing modified or locked nucleic acids (LNA) complementary to a specific gene or miRNA into cells without the use of traditional transfection reagents. Gymnosis has been used previously to knockdown gene expression in enteroids(34), however knockdown capacity specifically in IESCs has not been evaluated. To evaluate whether IESCs of *ex vivo* cultured enteroids would take up LNAs introduced through the media and/or matrigel via gymnosis and downregulate target gene/miRNA expression (Figure 4a), we tested knockdown efficacy of an LNA against EGFP (LNA-EGFP) in Lgr5-EGFP^+^ enteroids (Figure 4b, Supplemental Figure 5a-c). We identified EGFP^+^ crypts immediately after seeding into matrigel, and followed the growth of the enteroid over the course of 8 days (Supplemental Figure 5b). As Lgr5-EGFP crypts demonstrate mosaic expression (in our colony, approximately 1 in 30 crypts are EGFP^+^), qRT-PCR analysis was inconclusive (data not shown). However by Day 4, there was an appreciable depletion of EGFP based on fluorescence imaging, indicating successful knockdown of gene expression in IESCs of *ex vivo* enteroids using gymnosis.

**FIGURE 4.**
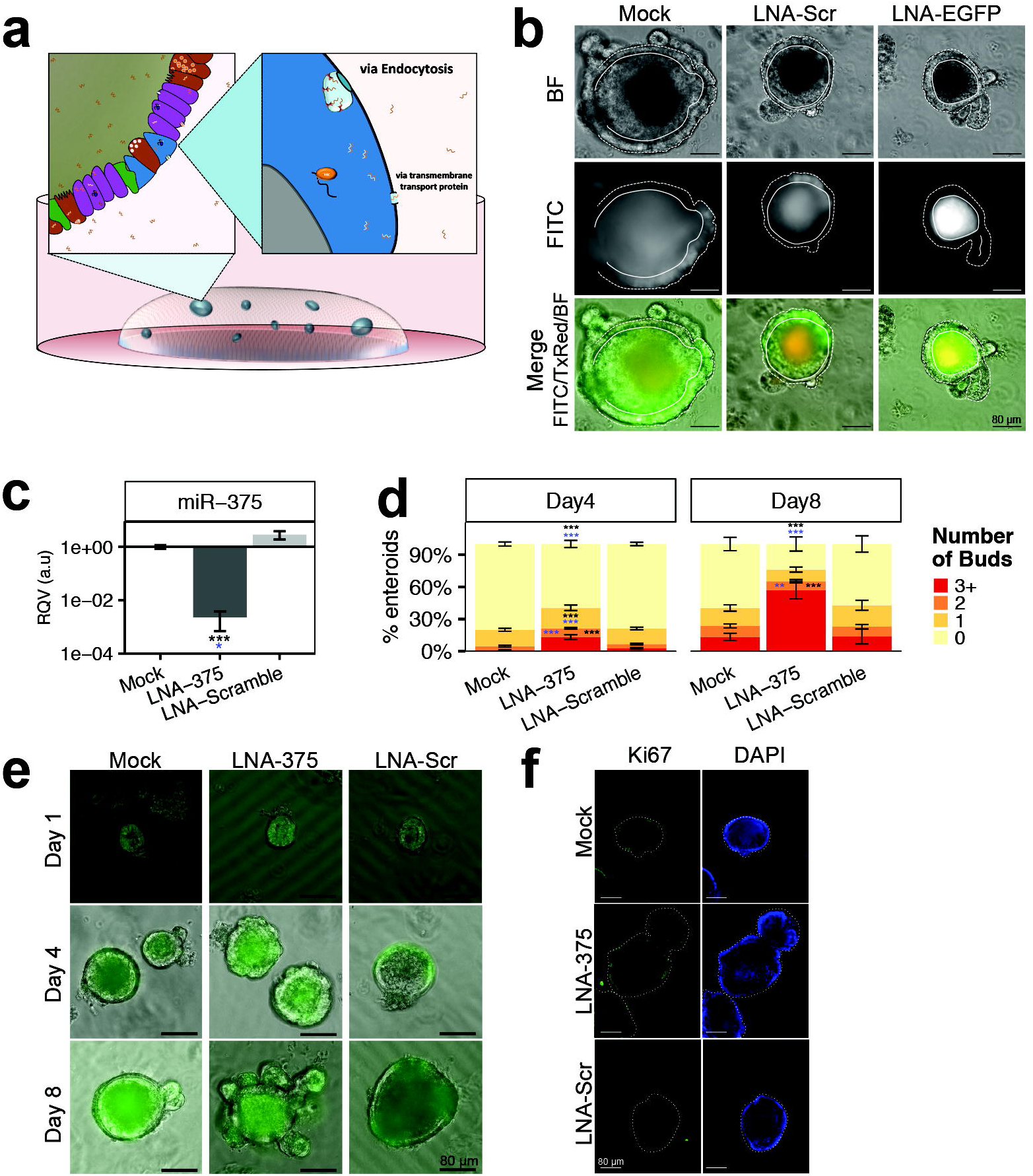
*Ex vivo* knockdown of miR-375 using gymnosis in enteroids results in increased proliferation. **a**) Schematic of miRNA knockdown in enteroids using gymnosis. **b**) Efficacy of gene knockdown in intestinal epithelial stem cells (IESCs) was evaluated by performing gymnosis of a locked nucleic acid (LNA) complementary to *EGFP* in Lgr5-EGFP+ positive enteroids. Lgr5-EGFP+ crypts were identified at Day 0 and tracked over the course of 8 days using fluorescence microscopy. A representative image of an EGFP+ enteroid is shown following gymnosis using no LNA (Mock), 1 μM scramble LNA (LNA-Scr), or 1 μM LNA complementary to EGFP (LNA-EGFP). Bright field (BF) and FITC fluorescence images are shown for each treatment. A final merged image contains stacked BF and FITC images, as well as TxRed, which was used to estimate autofluorescence. As enteroids are three-dimensional (3D) and filled with shed cells, the enteroid center may exhibit fluorescence due to its shape and density (autofluorescence) and/or accumulation of EGFP either secreted from or within shed cells. Dashed and solid white lines are used to indicate the region of focus on the 3D enteroid, which have minimal projection into the z-frame. Cells that fall between the solid and dashed lined are used to determine knockdown efficacy. **c**) Relative quantitative values (RQVs) are shown for miR-375-3p in Mock, LNA-375, and LNA-Scramble treated enteroids isolated from female germ-free (GF) Sox9-EGFP mice at Day 8 as measured by qRT-PCR relative to *U6*. **d**) Mean percent of enteroids isolated from female GF Sox9-EGFP mice with 0, 1, 2, or 3+ buds at Day 4 and Day 8 following Mock (n=12), LNA-375 (Day 4 n=12, Day 8 n=11), or LNA-Scramble (Day 4 n=12, Day 8 n=9) uptake by gymnosis. Data is combined from 3 independent experiments, consisting of 3-4 wells per condition. **e**) Representative images of enteroids isolated from female GF Sox9-EGFP mice at Day 1, Day 4, and Day 8, following Mock, LNA-375, or LNA-Scramble uptake by gymnosis. Sox9-EGFP expression (FITC/green) is overlaid on the bright field images. **f**) Confocal images of whole mount enteroids stained for Ki67 and DNA (DAPI). Experiments were performed in duplicate or triplicate. The ‘n’ refers to number of wells. Significance was determined using a Student’s two-tailed unpaired t-Test relative to Mock (black asterisks) or LNA-Scramble (blue asterisks). * p < 0.05, ** p < 0.01, *** p < 0.001. Error bars depict standard error of the mean. Scale bars depict 80 μm.

**FIGURE 5.**
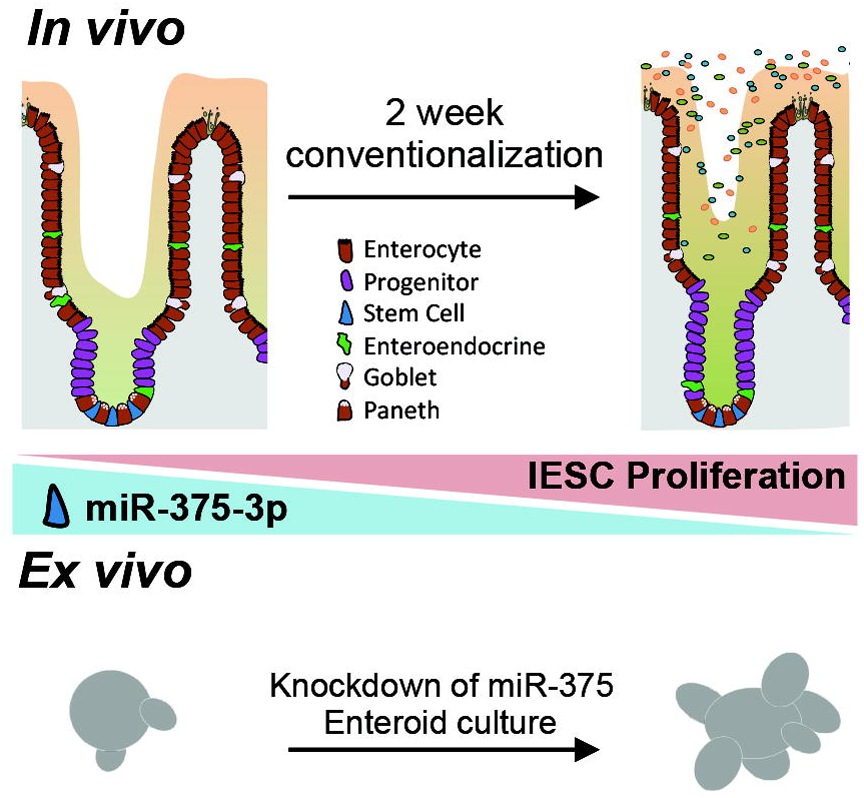
Current working model of miR-375-3p mediation of the effects of microbiota on intestinal epithelial stem cell (IESC) proliferation. Previous research shows increased intestinal epithelial proliferation upon conventionalization of germ-free (GF) mice. We found that miR-375-3p is down regulated in intestinal epithelial stem cells (IESC) upon conventionalization, and that *ex vivo* knockdown of miR-375-3p results in increased proliferative capacity.

*Knockdown of miR-375-3p in enteroids results in increased proliferation*—To test the functional effect of miR-375-3p downregulation, we knocked-down miR-375-3p by gymnosis in enteroids. We achieved a robust ~700-fold knockdown of miR-375-3p at day 8 using a LNA inhibitor complementary to miR-375-3p (LNA-375; Figure 4c). At both day 4 and 8, LNA-375 treated enteroids exhibited dramatically increased budding (Figure 4d & 4e, Supplemental Figure 6a-c), a marker of IESC proliferative capacity(35, 36), relative to Mock and LNA-Scramble treated enteroids. Consistent with this finding, whole mount staining of the enteroids also showed increased Ki67 upon knockdown of miR-375-3p (Figure 4f), though no difference in enteroid size (Supplemental Figure 6d) nor passage efficiency was observed (data not shown). These data indicate that miR-375-3p is a potent regulator of IESC proliferation and that microbiota may regulate IESC renewal in part via modulation of miR-375-3p (Figure 5).

## DISCUSSION

In this study, we have provided novel evidence that miRNAs are responsive to the presence of gut microbiota in a cell-type specific manner. Microbiota exert the strongest effect on host miRNA expression in the Sox9^Low^ population, which is highly enriched in IESCs(12-14, 16, 37). Subpopulation analysis was necessary to identify this effect, as IESCs make up only 1-3% of all IEC types. miR-375-3p was identified as significantly downregulated in the IESC population in response to microbiota, and follow-up experiments *ex vivo* demonstrated miR-375-mediated control of IESC expansion and proliferation, thereby providing a mechanism by which microbiota may regulate these processes during conventionalization *in vivo*. miR-375-3p has been associated previously with the regulation of proliferation and differentiation in several tissues(34, 38, 39). It is predicted to target many members of the Wnt/β-catenin and Hippo signaling pathways, but so far has only been experimentally shown to directly inhibit Frizzled-8(39) and Yap1(40). miR-375-3p has been knocked down systemically in mice, and while the authors did not study intestinal proliferation, they observed an increased rate of intestinal transit(41). miR-375-3p is best studied in the context of pancreatic endocrine cell differentiation and function(42-44), and more recently, Knudsen at al. (2015) identified a role for miR-375-3p in regulating EEC differentiation as well(34). We found that miR-375-3p is robustly expressed in both IESCs and EECs; however, we observed that miR-375-3p is responsive to microbiota only in IESCs (Figure 3d). This observation might suggest cell type specific microbial signaling pathways and cell type specific roles for miR-375-3p.

Our RNA-sequencing analysis, the first to our knowledge comparing GF and CV IESCs, demonstrates substantial gene expression differences in IESCs in response to microbiota. CV IESCs showed upregulation of genes associated with proliferation. Of note, our data indicate a ~4-fold increase in *Lgr5* mRNA expression (Supplemental Figure 3c). Microbiota may regulate Wnt signaling upstream of Lgr5, an R-spondin ligand receptor, as known upstream regulators of Lgr5 are also altered by the presence of microbiota, including *Gata4*, *Gata6*, and *Bmp4* (Supplemental Figure 3c)(45, 46). This is a finding that deserves further investigation.

An unexpected finding was that GF IESCs (Sox9^Low^) have a gene and miRNA expression profile demonstrating some similarity to Sox9^High^ cells. Given the careful sorting protocol, and the observation that Sox9^High^ enriched genes and miRNAs change in both the upward and downward directions within the Sox9^Low^ population, our finding is unlikely to be solely driven by contamination between populations. One possible explanation is that Sox9^Low^ cells are primed for the EEC lineage in the absence of microbial influence. Alternatively, one of the caveats of the Sox9-EGFP model is that while the Sox9^High^ populations consist primarily of EECs, they also include a small population of reserve stem cells(17). It is therefore possible that microbiota influence the maintenance of reserve stem cells in addition to their role in regulating actively cycling IESCs through miR-375-3p, though the gene expression data does not fully support this hypothesis. Though outside the scope of this study, more research including single cell analyses will need to be conducted to delineate more precisely the differences between GF and CV IESCs, as well as to determine which miRNAs are involved in the maintenance of active and quiescent IESC states.

An important added value of our study is the first ever map of miRNA expression across different IEC subtypes, and the cell type specific influence of microbial conventionalization on miRNA expression. We also provide evidence that IESC microbiota-sensitive miR-375-3p influences IEC proliferation, most likely through physiological maintenance of actively cycling IESC. Of course many questions still remain, including how microbiota influence miRNA expression in IESCs. This phenomenon may be explained by direct and/or indirect mechanisms. Regarding the former, although thus far bacteria have only been found to reside within the crypts of the caecum and colon, where microbial density is highest (47), it nevertheless remains a possibility that bacteria residing within the jejunal crypt may directly influence miRNAs in the stem cell subpopulation. Indirect mechanisms are also possible, such as changes in the microenvironment (metabolites and bacterial endotoxins) or through indirect signaling by immune or mesenchymal cells, which were not profiled in this study. Though outside the scope of this analysis, further research is certainly warranted to investigate the interesting relationship between host miRNAs and resident microbiota.

It is also important to note that each segment of the intestinal epithelium has distinct physiological roles and differing magnitudes of microbial load. Our study only examined changes in response to microbiota in IECs from the jejunum. In the future it would be important to assess differences in cell type specific responses to microbiota along the entire length of the intestine. Additionally, it would be interesting to investigate cell type specific responses to microbiota in other populations not sorted herein, including goblet and Paneth cell populations. These cell types do not express Sox9-EGFP, and are rare cell populations in the Sox9^Neg^ fraction, which is comprised primarily of enterocytes. Nevertheless, Paneth and goblet cells may experience robust changes in response to microbial presence based on their known functions. While our current study focuses on the Sox9-EGFP model, which precluded examining these populations, they deserve attention in future work.

From an experimental standpoint, our study also provided validation of an important new tool to knockdown gene expression in IESCs of intestinal enteroids using gymnosis, a technique that does not rely on cytotoxic transfection reagents(33, 34). While not fully investigated in our study, it is possible that gymnosis allows for the uptake of LNAs into other IEC types in addition to IESCs. A previous study using transfection of LNAs in an intestinal cell line, indicate stable knockdown of target miRNAs following a single transfection after 21 days(48), emphasizing the stability of LNAs in culture. While *ex vivo* culture demonstrates significantly more proliferation, further investigation of the stability of LNAs in enteroids, as well as the knockdown efficacy in other cell types is warranted to evaluate the full utility of this assay. Nevertheless, knockdown of gene expression in enteroids and in stem cells has proven quite difficult, requiring electroporation and adenoviral mediate knockdown. This study presents a quick and affordable alternative to knockdown gene and miRNA expression in IESCs of enteroids.

## CONCLUSIONS

In summary, we provide novel data on the miRNA landscape in four distinct cell populations from the intestinal epithelium, and demonstrate that miRNA profiles are very different across the IEC subtypes, and also that miRNAs respond to the presence of microbiota in a highly cell type specific manner. IESCs demonstrate robust gene and miRNA expression changes at 14 days post-conventionalization. We investigate one IESC microbiota-sensitive miRNA, miR-375-3p, and show that its downregulation results in significantly increased proliferative capacity, providing one possible mechanism by which microbiota regulate proliferation of IESCs *in vivo*. We believe the data provided herein progresses the field, and provides the scientific community a valuable resource through which researchers can initiate novel studies into miRNAs and microbiota-mediated regulation of intestinal physiology, homeostasis, and disease pathogenesis.

## EXPERIMENTAL PROCEDURES

*Animals*—All animal studies were approved by the University of North Carolina at Chapel Hill’s Institutional Animal Care and Use Committee. The original source and maintenance of *Sox9-EGFP* mice have been described elsewhere(12-14). Germ-free (GF) Sox9-EGFP mice on a C57BL/6J background were generated at the UNC Gnotobiotic Core Facility. Four pairs of female GF littermates were used in these experiments at 8-10 weeks of age. Each pair came from separate litters born between April and July 2015. GF mice were housed with animals of the same sex from the same litter, on Envigo 7070C Tekland Diamond Dry Cellulose bedding. Four age-matched conventionally-raised Sox9-EGFP animals and wild-type C56BL/6J animals were included as controls in each individual FACS experiment. Conventionally-raised and conventionalized mice were bedded on Andersons irradiated ¼. inch Bed-O’cobs laboratory animal bedding. The small RNA-seq data presented for conventionally-raised animals was generated in a separate experiment, which isolated each IEC population from female conventionally-raise animals fed a standard chow diet at 30-weeks of age. Crypt culture studies were performed using female conventionally-raised C56BL/6J, female GF Sox9-EGFP animals, and male conventionally-raised Lgr5-EGFP-IREScreERT2. Conventionally-raised animal colonies were maintained several generations at the University of North Carolina at Chapel Hill.

*Conventionalization*—For each littermate pair, 0.2-0.7 grams of fresh fecal pellets were collected on separate days from multiple animals across 6-8 cages in the conventionally raised Sox9-EGFP animal colony housed at UNC and were frozen at − 80C until reconstitution. Less than one hour before conventionalization, the fecal sample was thawed on ice and then reconstituted at 1 gram/10 mL cold PBS under anaerobic conditions. The fecal slurry was passed through a 100-μm filter to remove debris and 1 mL was aliquoted into a fresh microcentrifuge tube. For each littermate pair, one GF animal was conventionalized (CV) using prepared fecal slurry and administered by oral gavaged at 10 μL/gram body weight. To ensure conventionalization, whiskers and anus were swabbed and the remaining slurry was painted onto several pieces of food left on the bottom of the animal’s cage. CV animals were housed individually throughout the duration of conventionalization with access to food and water ad libitum.

*Intestinal epithelial cell (IEC) isolation and fluorescence-activated cell sorting (FACS)*—After a two-week conventionalization, both the CV and GF animals were anesthetized using isofluorane, then euthanized by cervical dislocation. The small intestine was removed and divided into 3 equal sections. The proximal and distal 10 cm were considered duodenum and ileum, respectively. The middle section was considered jejunum and used for all studies. Jejunum was flushed with ice cold PBS to remove contents, and total IEC were prepared for FACS as previously described(14). IECs were sorted using a Mo-Flo XDP cell sorter (Beckman-Coulter, Fullerton, CA) at the University of North Carolina Flow Cytometry Core Facility using previously described gating parameters(13-15). Conventionally-raised age-matched Sox9-EGFP animals were included in each individual sorting experiment and used to set Sox9-EGFP gates. CD31-APC (BioLegend, San Diego, CA), CD45-APC (BioLegend), and Annexin-V-APC (Life Technologies, Carlsbad, CA), and Sytox-Blue (Life Technologies) staining excluded immune cells, endothelial cells and apoptotic cells, respectively. Sox9-EGFP cells were then subsequently sorted based on Sox9-EGFP intensity directly into RNA lysis buffer (Norgen Biotek, Thorold, ON, Canada). Additionally, non-sorted IECs (NS) were collected for each animal, except one conventionalized mouse (CV314), which did not have enough remaining sample to isolate a NS IEC population. NS IECs were purified by FACS to exclude non-epithelial and dying cells, but were not sorted based on Sox9-EGFP intensity. Due to the density of cells, Sox9^Neg^ cells were sorted into cell culture media, then pelleted following sorting by centrifugation. Total RNA was isolated using either the Norgen Total RNA kit (Sox9^Neg^ & NS) or Norgen Single-Cell RNA Purification kit (Sox9^High^, Sox9^Low^, Sox9^Sublow^; Norgen) as per the manufacturer’s instructions. Nanodrop 2000 was used to quantify RNA.

*mRNA library preparation and sequencing*—mRNA-sequencing libraries were prepared from 10 ng total RNA using the Clonetech SMARTer Ultra Low Input library preparation kit combined with Nextera XT DNA sample preparation kit (Illumina) by the UNC High Throughput Sequencing Core Facility (as per the Clonetech sample preparation guide). Four libraries randomly pooled per lane and sequenced 100 bp single-end on a HiSeq2000 platform at the UNC High Throughput Sequencing Core Facility. Seven bases were trimmed from the beginning of each read using Trimmomatic (v0.36)(49) to eliminate remaining SMART adapter sequences, then reads were mapped and aligned to the GENCODE(50-52) mouse transcriptome (vM10) using STAR (v2.4.2a)(53) and SALMON (v0.6.0)(54). Transcript counts were then imported into R (v3.3.1), and were normalized and differential expression of genes quantified using DESeq2 (v1.12.4)(55, 56). Raw sequencing data as well as counts are available through GEO accession GSE81126.

*Small RNA library preparation and sequencing*—The small RNA sequencing was done at Genome Sequencing Facility of Greehey Children's Cancer Research Institute at University of Texas Health Science Center at San Antonio. Libraries were prepared using an average of 50 ng of total RNA using the TriLink CleanTag Small RNA Ligation kit (TriLink Biotechnologies, San Diego, CA) and suggested library preparation method. Six to seven libraries were pooled per lane, and were sequenced single-end 50x on the HiSeq2000 platform. One GF Sox9^Sublow^ sample failed during sequencing. However, for the remaining samples, we received an average of 26.5 million reads per sample. Raw sequencing data and miRNA quantification tables for all samples can be accessed through GEO record GSE81126.

*Bioinformatics*—Sequencing quality was extremely high as assessed using FASTQC. Reads were trimmed and aligned to the mouse genome (mm9) as previously described(29), with the following modification: only contigs with greater than one read alignment were passed into the to Shrimp alignment pipeline. An average of 58.9% of reads mapped to the mouse genome across samples (Mapping statistics can be found in Supplementary File 1). Due to the large number of reads mapping throughout the genome in GF315 NS IECs, Shrimp failed to align this sample, and it was eliminated from further analysis. Annotated miRNAs with a reads per million mapped to miRNAs (RPMMM) expression threshold of greater than 400 in at least one sample were used in further analyses. One aberrant CV Sox9^Sublow^ sample was identified on the basis of poor clustering by PCA and hierarchical clustering analyses, and was removed from subsequent analyses (Supplemental Figure 4).

*Enteroid culture*—Jejunum was isolated and flushed with cold PBS (Gibco cat. 14190-144, ThermoFisher Scientific, Waltham, MA), opened, and divided into 6 cm sections. Sections were placed in cold high glucose DMEM and rocked to remove excess fecal matter. Each section was then placed in 3 mM EDTA (cat 46-034-Cl, Corning, Corning, NY) diluted in PBS and rocked at 4ºC for 15 minutes. The luminal side of the tissue was gently scraped to remove villi and placed into fresh 3 mM EDTA/PBS and rocked an additional 30 minutes at 4ºC. Sections were shaken for 2 minutes in ice cold PBS to remove crypts, then filtered through a 70 μm cell strainer and counted. Crypts were resuspended into Reduced Growth Factor Matrigel (cat. 356230, BD Biosciences, Franklin Lakes, NJ). Advanced DMEM/F12 (Gibco, ThermoFisher) supplemented with GlutaMAX (Gibco cat. 35050-061, ThermoFisher), Pen/Strep (Gibco cat. 15140, ThermoFisher), HEPES (Gibco cat 15630-080, ThermoFisher), 1X N2 supplement (Gibco cat. 17502-048, ThermoFisher), 1 ng/uL EGF (cat. 2028-EG, R&D Systems, Minneapolis, MN), 2 ng/uL Noggin (cat # 250-38, PeproTech, Rocky Hill, NJ), 10 ng/uL murine R-spondin (cat # 3474-RS-050, R&D Systems), and Y27632 (cat. ALX-270-333-M025, Enzo Life Sciences, Farmingdale, NY) was added. For 48-well plates, 400 crypts were added to 10 μL matrigel, and 250 μL media was added to each well. For 96-well plates, 250 crypts were added to 5 μL matrigel, and 125 μL media added to each well. Either the media or matrigel was supplemented with PBS or an equal volume of LNA. For miRNA studies, LNAs were added at 500 nM and include miRCURY LNA Power Inhibitor against miR-375-3p (cat # 4101397, Exiqon, Woburn, MA) or Negative Control A (cat # 199006, Exiqon). For studies knocking down EGFP, a custom LNA longRNA Standard GapmeR was designed to target EGFP (Exiqon, Design ID: 590367-1), and Negative Control A Gapmer (Exiqon, cat# 300610) were used in concentrations ranging 500 nM – 5 μM. For EGFP knockdown studies, Lgr5-EGFP^+^ crypts were identified at Day 0 for follow up analyses. Enteroids were counted at Day 1 and bud formation assessed at Day 4 and Day 8 using an Olympus IX83 Inverted Microscope fixed with a live imaging incubator. Images of EGFP^+^ enteroids were take every 2 days. Media supplemented with half the original starting concentration of LNA or PBS when changed at Day 4, and growth factors were supplemented every other day. Enteroids were harvested at Day 8 and RNA was isolated using the Norgen Total RNA isolation kits as per manufacturers instructions.

*Whole mount staining of enteroids*—For whole mount staining, enteroids were fixed in 2% PFA, then permeabilized using 0.5% Triton X-100 (Sigma-Aldrich) diluted in PBS. Enteroids were blocked using 10% Normal Goat Serum diluted in our immunofluorescence buffer, which consisted of 0.1% bovine serum albumin (Sigma-Aldrich), 0.2% Triton X-100, and 0.05% Tween-20 (Sigma-Aldrich) in PBS. Then, enteroids were stained using antibodies against and Ki67 (ab15580, 1:250, Abcam, Cambridge, MA) diluted in immunofluorescence buffer. Nuclear staining was done using Hoechst 33342 (1:5000, cat. H3570, ThermoFisher) diluted in immunofluorescence buffer. Confocal imaging was performed at the UNC Microscopy Core Facility on a Zeiss CLSM 710 Spectral Confocal Laser Scanning Microscope.

*Analysis of enteroid area*—At day 8, a z-stack of 4x bright field images capturing the entire well was taken every 10 μm throughout the matrigel patty. Each enteroid was measured for area at its maximal projection within the z-stack as an estimation of enteroid size, using imageJ. All enteroids that reached at least 1000 μm^2^ within each well were included. Three wells from each condition were analyzed from a single experiment.

*Validation of miRNA expression levels*—miRNA expression in the CV and GF animals was validated by qRT-PCR using Taqman assays (Applied Biosystems, Foster City, CA). Relative quantitative value (RQV) was determined relative to control gene *U6*.

*Linear Model*—The model covariates include cell type, *T*; condition, *C*; littermate pair, *P*; and sequencing group, *G*; as well as an interaction term between cell type and condition (Equation 1).

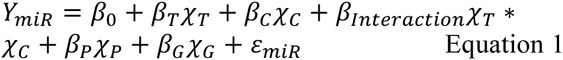

To determine significance, a multiple testing correction (False Discovery Rate) was performed on p-values for each covariate across all miRNAs.

## Acknowledgements

Many thanks to R. Eric Blue for help in developing the GF Sox9-EGFP resource; Elaine Glenny for critical assistance with the anaerobic chamber and the protocol for conventionalization; Dr. C. Lisa Kurtz, Dr. Emily Moorefield, Dr. Jeanette Baran-Gale, and Dr. Kasia Kedziora for technical assistance and training; Dr. John Rawls and Dr. Scott Magness for helpful discussions; as well as Felicia Heyward and UNC Flow Cytometry Core Facility, the UNC Gnotobiotic Core Facility, Dr. Zhao Lai and the UTHSCSA Genome Sequencing Facility, and Dr. Bob Bagnell and the UNC Microscopy Services Laboratory for critical services.

## Author contributions

BCEP conceptualized experiments, conducted experiments, analyzed and interpreted data, prepared figures, drafted and revised the manuscript. AM worked with the UNC Gnotobiotic core facility to develop the GF Sox9-EGF model. WP conducted experiments, analyzed and interpreted data. SD assisted with experimental design, and provided technical advice, and assistance with data interpretation. PKL provided assistance with conceptualization and experimental design, and contributed to the revision of the manuscript. PS obtained funding, provided assistance with conceptualization and experimental design, interpreted data, supervised the study, and revised the manuscript.

## FOOTNOTES

Research was supported by UNC GMB T32GM007092 (B.P., NIGMS/NIH via UNC Curriculum in Genetics & Molecular Biology), UNC IMSD Diversity Fellowship (B.P. via 5R25GM55336, NIGMS/NIH), F31DK105747 (B.P., NIDDK/NIH), 14CSA20660001 (P.S., American Heart Association), P30DK034987 (P.S., NIDDK/NIH via UNC Center for Gastrointestinal Biology and Disease), 1-16-ACE-47 (P.S., American Diabetes Association), and R01DK105965 (P.S., NIDDK/NIH). The UNC Gnotobiotic Core Facility is supported through NIH grants P39DK034987 and P40OD010995.

The abbreviations used are: IESC, intestinal epithelial stem cell; microRNA, miRNA; IEC, intestinal epithelial cell; EEC, enteroendocrine cell; FACS, fluorescence-activated cell sorting; GF, germ-free; CV, conventionalized; EGFP, enhanced green fluorescent protein; CR, conventionally-raised; NS, non-sorted; RPMMM, reads per million mapped to miRNAs; PCA, principal component analysis; LNA, locked nucleic acid; FC, fold-change; FDR, false discovery rate.

## SUPPLEMENTAL DATA LEGENDS

**SUPPLEMENTAL FIGURE 1.** Physiological parameters observed at the time of sac of female germfree (GF) and 2-week conventionalized (CV) mice at10-12 weeks of age (n=4 each). **a**) Weight of spleen in grams. **b**) Length of small intestine (SI) from pyloric sphincter to caecum in centimeters (cm). **c**) Length of the colon from caecum to anus in cm. **d**) Ratio of spleen weight to body weight. **e**) Body weight in grams. **f**) Liver weight in grams. **g**) Ratio of liver weight to body weight. Significance was determined using a Student’s two-tailed paired t-Test and is denoted as follows: * p < 0.05.

**SUPPLEMENTAL FIGURE 2**. Example gating scheme for fluorescence-activated cell sorting (FACS) of Sox9-EGFP intestinal epithelial cells (IECs). **a**) Cells were gated to remove dead cells, as well as cellular debris using side-scatter (SSC) height by forward scatter (FSC) area. **b** & **c**) Then, singlets were selected for by gating on SSC-Height by SSC-Area and then FSC-Width. **d**) Dying and immune cells were gated out using Sytox blue staining, and CD31-APC, CD45-APC, and Annexin V-APC staining, leaving a highly pure IEC population with which to gate on EGFP intensity. **e**) Distinct subpopulations of IECs were isolated based on their cellular EGFP intensity. Gating for germ-free and conventionalized animals were set using a conventionally-raised Sox9-EGFP animal, representative sort images are from sort of animal CV314.

**SUPPLEMENTAL FIGURE 3**. RNA-seq of the Sox9^Low^ population from germ-free and conventionalized animals. **a**) Volcano plot showing differentially expressed genes in Sox9^Low^ intestinal epithelial stem cells (IESCs) between germ-free (GF) and conventionalized (CV) mice. Horizontal dashed grey line indicates a False Discovery Rate (FDR) of 0.05. Vertical dashed grey lines indicate fold change (FC) > 1.5 or < −1.5. Significantly changed genes are colored in orange and red, representing genes that at enriched in CV or GF IESCs, respectively. **b**) Top Enrichr Gene Ontology Biological Process enrichment terms for genes significantly upregulated in GF (left) or CV (right) IESCs. **c**) Relative quantitative values (RQV), which is in arbitrary units (a.u), of normalized counts for selected genes in CV and GF IESCs (n=4 animals per condition). Genes selected include *Sox9*, those known as markers for enteroendocrine (EEC) cell types, other differentiated lineages, including Paneth cells (*Lyz*), goblet cells (*Muc2*), enterocytes (*Elf3*), reserve/quiescent IESCs (rIESCs), actively cycling IESCs (aIESCs), transcriptional regulators of *Lgr5* that promote proliferative potential, as well as other markers of proliferation. Significance was determined using DEseq2 differential expression analyses, combined with multiple testing correction, and is denoted as follows: * FDR < 0.05, ** FDR < 0.01, *** FDR < 0.001. Error bars depict standard error of the mean.

**SUPPLEMENTAL FIGURE 4**. Principal components analysis (PCA) applied to miRNA expression in reads per million mapped to miRNAs (RPMMM) for all miRNAs that reached an expression threshold of at least 400 in 1+ samples (n=149). Biplots of variance projections of PC1 on PC2 and PC1 on PC3 are shown. Percent variance explained by each component is shown in parentheses along each axis. Samples are colored based on the cell population of origin, and the shape of the point indicates whether the sample came from a conventionally-raised (CR), germ-free (GF) or conventionalized (CV) mice. Circled in red is Sample CV246_Sublow, which was removed from subsequent analyses due to poor clustering.

**SUPPLEMENTAL FIGURE 5**. Gymnosis is effective at knocking down gene expression in Lgr5-EGFP^+^ stem cells. Approximately 1 in 30 crypts were positive for EGFP. To evaluate knockdown, positive crypts were identified immediately after seeding in 96 well plates, and were followed for 8 days. **a**) Media was supplemented with various concentrations of locked nucleic acids (LNAs) complementary to *EGFP* (LNA-EGFP), a scramble control (LNA-SCR), or no LNAs (Mock). At day 8, media was changed and 50% of the original concentration of LNA was supplemented. An example of an EGFP^−^ enteroid is included from a Mock treated well. Top corner inset shows Day 0 enteroids with positive EGFP expression. **b**) Time course showing growth of an EGFP^+^ enteroid treated with 1 μM LNA-SCR or 1 μM LNA-EGFP. **c**) EGFP^+^ enteroids treated with 1 μM LNA-SCR or LNA-EGFP are shown. To demonstrate within-well variability, all EGFP^+^ enteroids from one well of LNA-EGFP treated enteroids are shown at day 8. EGFP^−^ enteroids are marked with a red ‘x.’ (**a-c**) The images within contain bright field and/or FITC and TxRed stacked images. As enteroids are three-dimensional (3D) and filled with shed cells, the enteroid center may fluorescence due to on its shape and density (autofluorescence) and/or accumulation of EGFP either secreted from or within shed cells. TxRed was used to estimate autofluorescence in 3D enteroids. Dashed and solid white lines are used to indicate the region of focus on the 3D enteroid, which have minimal projection into the z-frame. Cells that fall between the solid and dashed lined are used to determine knockdown efficacy. Scale bars depict 80 μm.

**SUPPLEMENTAL FIGURE 6**. Knockdown of miR-375 in enteroids isolated from several mouse models results in increased budding. Mean percent of enteroids with 0, 1, 2, or 3+ buds at Day 4 and Day 8 following treatment with no locked nucleic acids (LNAs, Mock), LNA complementary to miR-375 (LNA-375), or a scramble control (LNA-Scramble) are shown for: **a**) Enteroids generated from male Lgr5-EGFP mice (Mock, n=7; LNA-375 n=6; LNA-Scramble, n=6). **b**) Enteroids generated from female C57BL/6J mice (Mock, n=16; LNA-375, n=16; LNA-Scramble, n=12). **c**) Enteroids generated from female germ-free Sox9-EGFP mice (Mock, n=12; LNA-375, n=11; LNA-Scramble, n=9). Experiments were performed in duplicate or triplicate. The ‘n’ refers to number of wells. Significance was determined using a Student’s two-tailed unpaired t-Test relative to Mock (black asterisks) or LNA-Scramble (blue asterisks). * p < 0.05, ** p < 0.01, *** p < 0.001. Error bars depict standard error of the mean. **d**) Distribution of areas of the maximal projection of all enteroids isolated from male Lgr5-EGFP mice after 8 days following Mock treatment, 1 μM LNA-375, or a 1 μM LNA-Scramble. (n=3 wells each). **e**) Representative 4x images of wells treated with Mock, 1 μM LNA-375, or a 1 μM LNA-Scramble quantified in panel **d**.

**SUPPLEMENTAL FILE 1.** Linear regression model reveals differential expression of miRNAs across cell type, condition, sequencing batch and littermate pairs. This table contains false discovery rate (FDR) values for each regression covariate for all miRNAs that reached an expression threshold of at least 400 in 1+ sorted samples (n=149).

